# Deep, Staged Transcriptomic Resources for the Novel Coleopteran Models *Atrachya menetriesi* and *Callosobruchus maculatus*

**DOI:** 10.1101/035998

**Authors:** Matthew A. Benton, Nathan J. Kenny, Kai H. Conrads, Siegfried Roth, Jeremy A. Lynch

**Affiliations:** Institute for Developmental Biology, University of Cologne, Zülpicherstrasse 47b, Cologne, 50674, Germany; Simon F.S. Li Marine Science Laboratory of School of Life Sciences and Center for Soybean Research of the State Key Laboratory of Agrobiotechnology, The Chinese University of Hong Kong, Shatin, Hong Kong; Department of Biological Sciences, University of Illinois at Chicago, Chicago, Illinois, United States

**Keywords:** Beetle, Coleoptera, Transcriptome, Ovary, Embryo, Staged, Atrachya menetriesi, Callosobruchus maculatus, Chrysomelidae

## Abstract

Despite recent efforts to sample broadly across metazoan and insect diversity, current sequence resources in the Coleoptera do not adequately describe the diversity of the clade. Here we present deep, staged transcriptomic data for two coleopteran species, *Atrachya menetriesi* (Faldermann 1835) and *Callosobruchus maculatus* (Fabricius 1775). Our sampling covered key stages in ovary and early embryonic development in each species. We utilized this data to build combined assemblies for each species which were then analysed in detail. The combined *A. menetriesi* assembly consists of 228,096 contigs with an N50 of 1,598 bp, while the combined *C. maculatus* assembly consists of 128,837 contigs with an N50 of 2,263 bp. For these assemblies, 34.6% and 32.4% of contigs were identified using Blast2GO, and 97% and 98.3% of the BUSCO set of metazoan orthologs were present, respectively. We also carried out manual annotation of developmental signalling pathways and found that nearly all expected genes were present in each transcriptome. Our analyses show that both transcriptomes are of high quality. Lastly, we performed read mapping utilising our timed, stage specific RNA samples to identify differentially expressed contigs. The resources presented here will provide a firm basis for a variety of experimentation, both in developmental biology and in comparative genomic studies.

## Introduction

The order Coleoptera is the most speciose clade of animals currently known. Despite the best efforts of generations of biologists, its species are only sparsely sampled and are yet to be comprehensively described, with approximately 90% of coleopteran diversity as yet uncategorized (e.g. [1,2]). A similar discrepancy exists at the molecular level; while several genomic resources are available in this clade, their number and phylogenetic distribution is only just beginning to accurately sample that of the Coleoptera as a whole. A wide range of transcriptomic data is available in whole organisms (for example [3–5], among others), specific body parts (such as [6,7]), and in several cases for staged embryos following RNA interference-mediated gene knock-down [8–10]. The i5K project [11] will also greatly advance our knowledge of the genomic complement of Coleopterans, with 69 species of this Order listed on that database as nominated for genomic sequencing as of the 07/04/16 (url: http://arthropodgenomes.org/wiki/i5K_nominations) and several genomes publically available [12–14]. However, the majority of this information is largely still outside of the public domain, the Coleoptera are still relatively undersampled compared to the Diptera and Hymenoptera, and, in particular, timed embryonic resources (e.g. [15]) are rare in the literature.

The true phylogeny of the Coleoptera is still under investigation but, in general, four suborders, 17 superfamilies, and 168 families are recognised [16]. The structure of the coleopteran clade can be seen summarised in Fig 1A. Coleopterans have long been used for research into embryology, and in the pre-molecular era, the Chrysomelidae (summarised phylogeny shown in Fig 1B) was one of the best studied superfamilies. For example, the first functional embryonic experiments in any insect were carried out in the Colorado potato beetle, *Leptinotarsa decemlineata* (sub-family Chrysomelinae, see Fig 1B), leading to the discovery of the function of pole cells and the existence of germ plasm in insects [17,18]. Further, the larval cuticle preparation technique, so vital for arthropod developmental biology, was first perfected in the bean beetle, *Callosobruchus maculatus* [19]. Chrysomelid beetles are also interesting from an ecological and economic viewpoint, as members of the group are usually pest species, perhaps most famously the aforementioned Colorado potato beetle, which is a major pest of potato crops in America, Asia and Europe [20].

**Figure 1:**
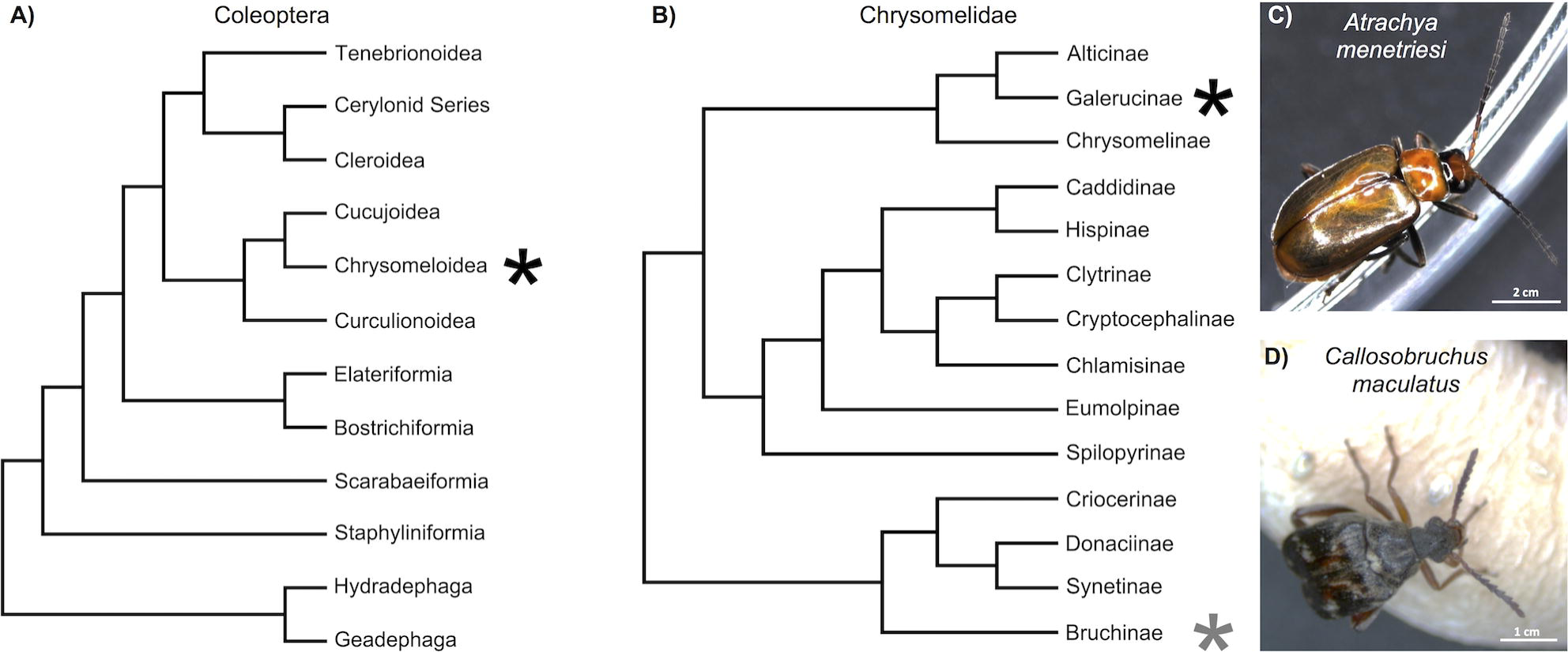
A) Cladogram of Coleoptera simplified from that determined by [16]. Black asterisk indicates the superfamily in which the Chrysomelidae are located. B) Cladogram of Chrysomelidae simplified from that determined by [21]. Black and grey asterisks indicate the sub-families in which *A. menetriesi* and *C. maculatus* are located (respectively). C) *A. menetriesi* and D) *C. maculatus* adults.

The Chrysomelidae are represented in public databases by a number of ongoing transcriptomic and genomic projects. In particular, both *L. decemlineata* (Bioproject PRJNA171749) and *C. maculatus* [22] are the subject of ongoing genomic sequencing. However, while a number of transcriptomes are planned or published in this clade (e.g. [5,6,23–27]) none yet sample across embryonic time points in a fashion which would allow insight into the genetic mechanisms behind key developmental stages or deep sampling of developmentally important genes. Here we present transcriptomic sequences for two species of chrysomelid beetle, the false melon beetle, *Atrachya menetriesi,* and the aforementioned *C. maculatus* (each of which are pictured along with their relative phylogenetic positions in Fig 1).

Compared to *C. maculatus,* the false melon beetle *A. menetriesi* (Faldermann), is a relatively unknown and understudied species. It is native to Japan, where it is an agricultural pest, and feeds on a variety of plants such as clover and lettuce. Although there has only been a small amount of research carried out on this beetle, the work that has been done has highlighted several interesting developmental traits, including the possibility of generating twin embryos after egg bisection [28], or up to four, seemingly complete, embryos following treatment with low temperatures [28,29]. Another interesting trait exhibited by this beetle is that almost all eggs enter diapause at a certain stage, and this is only broken in the wild by winter conditions. However, a small proportion of eggs skip this diapause and continue to adulthood. The ratio of diapause to non-diapause eggs varies in different parts of Japan [30], and is a heritable trait [31]. Further, *A. menetriesi* embryos undergo a very short germ band mode of development [32], contrasting strongly with beetles such as *C. maculatus* (see below). As the last common ancestor of these two species is estimated to have existed only 80 million years ago [21], how their developmental mechanisms have diverged so greatly is a potentially fascinating area for future study. Research on these topics would greatly benefit from modern molecular studies.

*C. maculatus* (Fabricius) is native to West Africa [33] but is now found worldwide, and is a common pest of stored legumes. It is also known as the southern cowpea weevil, however, it is not a true weevil. As noted above, this beetle was the focus of active developmental research in the pre-molecular era, with special focus on segmentation [34,35]. Segmentation in *C. maculatus* has also been studied more recently via immunohistochemistry for the even-skipped protein [36]. This recent work confirmed previous reports that the embryos undergo the long germ mode of development, similar to dipterans like *Drosophila melanogaster* and hymenopterans like *Nasonia vitripennis* [37]. It is commonly believed that the long germ mode of development evolved independently in dipterans and hymenopterans, and given the phylogenetic distribution of short and intermediate germ development within the Coleoptera [38,39], it seems likely that the long germ mode of development also evolved independently in the clade to which *C. maculatus* belongs. A comparison of the molecular basis of development in *C. maculatus* and other long germ insects, plus with more closely related species that feature short germ development, such as well studied flour beetle *Tribolium castaneum* (super-family Tenebrionoidea, see Fig 1A) and *A. menetriesi*, would yield crucial information on how developmental pathways have evolved to generate the long germ mode of development. *C. maculatus* has also been studied in other fields and is a useful system for undergraduate lab teaching [22] which could be aided further with deeper sequence resources.

In order to facilitate research on *A. menetriesi* and *C. maculatus*, as well as wider investigations in the Coleoptera and beyond, we present here deep, multi-stage transcriptomic resources from a range of key time points in the development of these two beetle species. Using RSEM-based methods we have compared transcript abundance across these life stages, which will allow the investigation of genes that play key roles in developmental changes in these species, particularly at the maternal-zygotic transition. We have carried out extensive searches for key genes in developmental patterning and cell signalling pathways, and from our analyses we conclude that the transcriptomes for both species are of very high coverage, with almost all expected genes being present with long (likely full) average open reading frames. We have already made extensive use of these resources for our own studies on the embryonic development of these two beetles, and are confident that they will be of broad utility to a range of fields in genomics and developmental biology.

## Materials and Methods

### Animal Husbandry

*A. menetriesi* eggs were collected from the wild and kindly provided by Dr Yoshikazu Ando, and were reared at 25°C on wet sand or soil and fed fresh lettuce. *C. maculatus* beetles were kindly provided by Dr Joel Savard, and were reared at 30°C on dry black-eyed peas.

### RNA Extraction and Sequencing

RNA was extracted from dissected ovaries and timed embryonic samples using a TRIzol RNA extraction kit according to the manufacturer’s protocols. RNA quantity and quality was checked using a Thermo Scientific Nanodrop 2000C Spectrophotometer and 1 µg was sent for sequencing by the Cologne Centre for Genomics on a Illumina HiSeq 2000 sequencer after sample preparation using a TruSeq RNA Library Preparation Kit (Illumina). Adaptor trimming and initial quality control was performed by the provider according to their proprietary standards, with no orphan reads kept. This cleaned data was then made available to us for download from an external server. Paired end read quality after sequencing was assessed using the FastQC program [40] and no residual adaptor was observed, as detailed in the Results.

### Transcriptome Assembly and Comparative Expression Analyses

Assemblies used in our final analysis were made using Trinity version 2013_08_14 [43], with the default settings (--min_contig_length 200). Trimmomatic [41] was assayed (Illuminaclip Leading:3 Trailing:3 Slidingwindow:4:15 Minlen:36) but not utilised for assemblies presented here, as described in results. Full assemblies were made using reads from all time points, and individual assemblies were then constructed using reads from each sampled time point individually. All assemblies are available from Figshare online (*A. menetriesi* DOI: 10.6084/m9.figshare.2056464.v2, *C. maculatus* DOI: 10.6084/m9.figshare.2056467.v2). DeconSeq standalone version 0.4.3 [44] was run on full assemblies with settings -i 95 -c 95, using the bact, fungi, hsref, and prot databases. Comparative expression analyses were performed using RSEM [61] as packaged in the Trinity module (cross sample normalisation: Trimmed Mean of M-values), to compare staged RNA samples with the combined assembly. Results shown here are the ‘as-isoform’ data, although ‘as-gene’ data is also provided in Additional Files. BUSCO v1.1b1 [48] was used to assess gene complement completeness.

### Functional Annotation

Our combined assemblies were automatically assigned homologs and annotated according to gene ontology (GO) terms using Blast2GO [49,50]. Initially, BLASTx was run using BLAST 2.2.29+ against the NCBI nr database as downloaded to a local server on the 17/01/2015, with settings -evalue 0.001 -max_target_seqs 5 -outfmt 5. GO term distribution within the *D. melanogaster* and *H. sapiens* genomes were downloaded from B2GO-FAR [62] and calculated using the Combined Graph function of Blast2GO. KEGG KAAS mapping was automatically performed using the KEGG KAAS tool (http://www.genome.jp/tools/kaas/), single-directional best hit with default BLAST settings, and with the eukaryote dataset as a basis for annotation.

### Gene Identification

Gene sequences were manually identified and their homology confirmed by independently using tBLASTn [63] searches using gene sequences of known homology downloaded from the NCBI nr database as queries against standalone databases created on a local server using BLAST 2.2.29+ or the CLC Main workbench. Genes putatively identified using this method were reciprocally BLASTed against the online NCBI nr database using BLASTx to confirm their identity. Where identity was uncertain, phylogenetic analysis was used to confirm identity.

### Phylogenetic Tree Construction

Sequences were aligned using MAFFT 7 [64] unless otherwise stated under the L-INS-i strategy. Alignments were then saved and exported to MEGA 6, where regions of poor alignment were manually excluded and maximum likelihood phylogenetic trees were constructed using the LG model, 1000 bootstrap replicates as indicated, 4 gamma categories and invariant sites, and all other default prior settings [65].

## Results and Discussion

### RNA Extraction and Sample Selection

Ovaries and embryos were collected as described in Materials and Methods, and as seen in Fig 2. The chosen time-windows cover a variety of important stages in the development of these species, and when combined can reasonably be expected to contain the majority of embryonic transcripts in their expressed complement. Briefly, RNA was collected from dissected ovaries from each species and from four embryonic time windows for *A. menetriesi* and two embryonic time windows for *C. maculatus*. Samples were sequenced on the Illumina HiSeq 2000 platform (one lane per species), adaptor trimming was performed, along with preliminary assessment of read quality, and data was made available for download from an external server.

**Figure 2:**
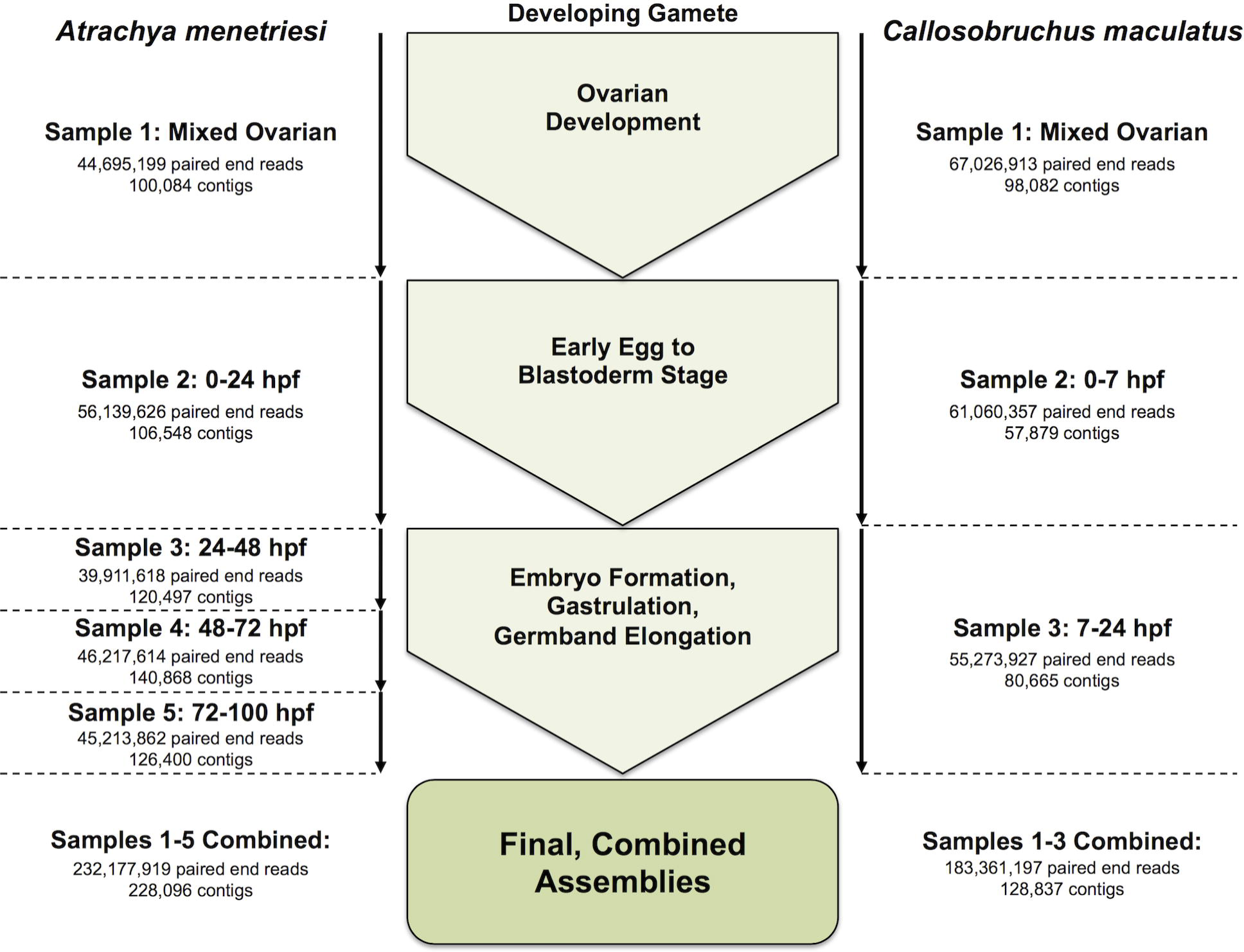
Summary of RNA sources, life stages sampled and sequencing results, with *A.menetriesi* data presented at left and *C. maculatus* data at right.

### Read Quality and Assembly Metrics

Fig 2 summarises initial read information, and these reads are available from the NCBI SRA, Bioproject Accession numbers: PRJNA293391 and PRJNA293393. To confirm quality of read data, FastQC [40] was run on all read data. This showed Phred quality scores were high, with median scores always exceeding 28 through to the 101st base, and generally in the mid-30s. A slight bias was found in initial nucleotide sequence. To ensure that Illumina adaptors were removed in their entirety, Trimmomatic v.0.32 [41] was used to check for residual adaptor sequence, but none were observed. We therefore posit that this bias is due to known biases in Illumina hexamer binding [42] rather than sequencing-based artifacts.

Using Trinity [43] as an assembler, Trimmomatic-corrected read assemblies were then compared with assemblies based on uncorrected reads. Initial assays of Trimmomatic-treated read assemblies empirically found them to be less well assembled than those using un-trimmed reads, with a shorter N50 and less total sequence recovery (total number of bp, of which some contigs could result from read errors). Given the advantages of a longer N50 and a preference to preserve as much information as possible, trimmed assemblies were therefore discarded in favour of untrimmed assemblies given the reasonable coverage requirement, and these ‘untrimmed’ assemblies were used for all further analyses.

Assays of initial assemblies noted a small amount of fungal contamination in the data for both species. This contamination likely comes from the environments in which the beetles are cultured. To correct for this, DeconSeq [44] was run on the Trinity output for both species, with high stringency as noted in the methods. A total of 336 and 206 contigs with some homology to fungal sequence was removed from the *A. menetriesi* and *C. maculatus* total assemblies as a result, before read mapping was performed. In the *A. menetriesi* assembly, we observed particularly high contamination with the protists, notably *Dictyostelium discoideum* and *Naegleria gruberi,* and some of this almost certainly remains in the transcriptome, as is normal with most ‘omics’ experiments. Metrics for final assemblies after the removal of contamination can be seen in Table 1, alongside the results of assemblies for individual time points.

**Table 1:**
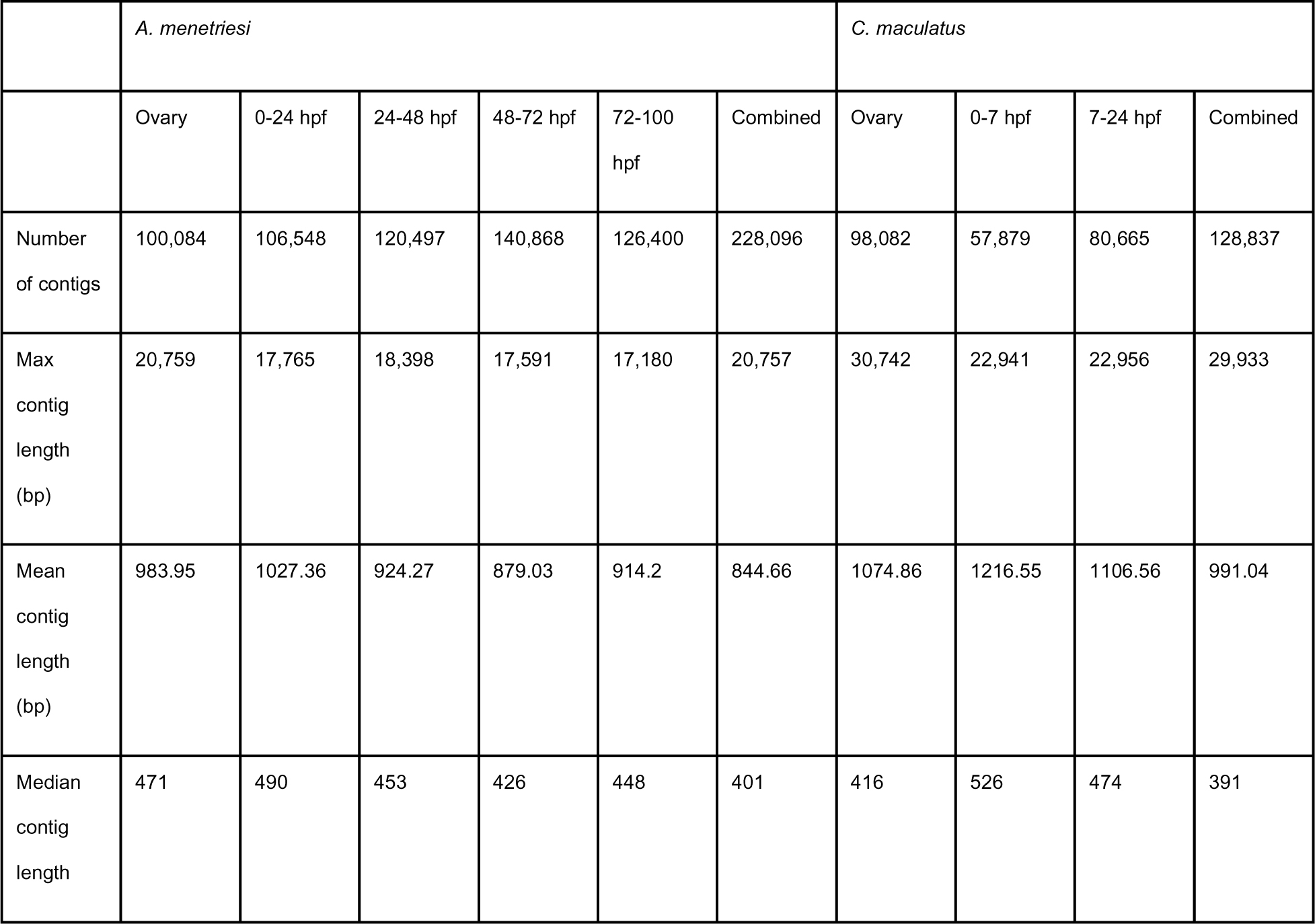
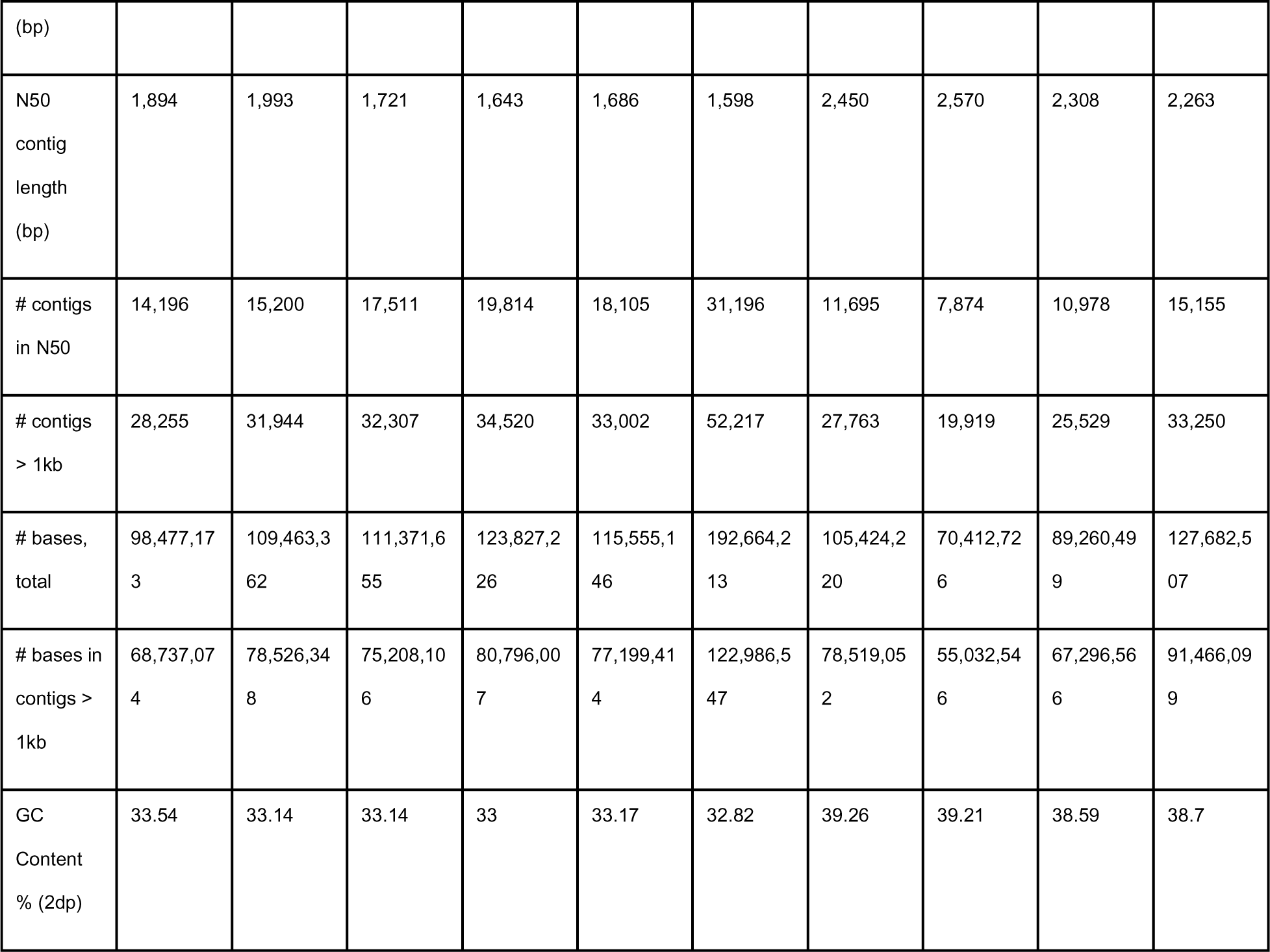
Assembly metrics for individual time point and combined transcriptomic resources

The final, combined timepoint, contamination-removed assemblies, comprising 228,096 and 128,837 contigs for *A. menetriesi* and *C. maculatus* respectively, contain a large number of well-assembled transcripts, with the number of contigs greater than 1 kb in length (52,217 in *A. menetriesi,* 33,250 in *C. maculatus*) and a high N50 (1,598 bp *A. menetriesi,* 2,263 bp *C. maculatus*) indicating a very well assembled dataset. This size is sufficient to span most protein coding domains, allowing easy inference of homology, and will be the full length of many transcripts. It is important to note that many of the contigs in our assembly will represent splice variants of single genes, and some genes will have multiple splice variants, which will affect these statistics. However, excellent recovery of splicing variation itself will be useful for a range of later analysis. GC content of the final assembled transcriptomes closely mirrors that of reads (32.82% in *A. menetriesi,* 38.7% in *C. maculatus*; reads 36-38%/40-43%, respectively, depending on library), and is therefore similar to that expected.

To gain an understanding of whether full-length transcripts were present in our data, we ran TransDecoder v 2.01 [45] to identify open reading frames (ORFs) and filtered for the results that were at least 100 amino acids long. For the *A. menetriesi* assembly, this analysis yielded 71,961 raw and 51,912 filtered contigs, while for the *C. maculatus* assembly, the analysis yielded 65,433 raw and 36,535 filtered contigs. The mean average length of the predicted polypeptides, after the filtering step, was 351 (*A. menetriesi*) and 448 (*C. maculatus*) amino acids. This is long enough for us to be confident that our assembly adequately spans coding regions, as the average eukaryotic protein length is 361 amino acids [46]. Together, these analyses suggest that we have recovered the vast majority of coding sequence in our combined assemblies, with sufficient length to adequately span ORFs, a conclusion further supported by gene annotation data as discussed further below. The numbers of ORFs presented here are considerably more than the 16,404 gene models observed in the *T. castaneum* genome (*Tribolium* Genome Sequencing Consortium, 2008), and this is likely due to both spurious ORFs in our dataset and to multiple splice variants.

Further information was gained using the Ortholog Hit Ratio method detailed in [47], which describes the proportion of best-hit ortholog sequence represented by a dataset. When compared to the *T. castaneum* Tcas3.31.pep.all.fa peptide set with a BlastX cut off of *E*^-5^, the average ratio of the full length ortholog present in all of our blast hits was 0.4369 (*A. menetriesi*) and 0.5079 (*C. maculatus*). These statistics compare very well with those found in other organisms and previous studies, such as that in [47], and further indicate that the *C. maculatus* transcriptome may be slightly better assembled than the *A. menetriesi* dataset. This difference is likely due to a major difference in the level of genetic heterogeneity between our samples from the two species, as *C. maculatus* have been cultured in our lab for many years, while *A. menetriesi* was recently sourced from the wild.

### Timed RNA Expression - Differential Expression Analysis

Key developmental stages for both *A. menetriesi* and *C. maculatus* can be easily observed following fixation and nuclei staining (data not shown). Briefly, egg lay to uniform blastoderm stage takes approximately 24 hours in *A. menetriesi*, and 7 hours in *C. maculatus*. Germband formation, gastrulation and elongation occur from 24-100 hours in *A. menetriesi*, and from 7-24 hours in *C. maculatus*. The latter period was subdivided in *A. menetriesi* according to characteristic stages of short germ development pertinent to our research interests. These stages, along with assayed sample periods, are shown diagrammatically in Fig 2. As well as being used for combined assemblies as described earlier, RNA extracted from mature ovaries and from embryos collected during the aforementioned time periods was also individually assembled using Trinity, allowing this information to be used to find time/stage-specific transcripts within our dataset.

The timed transcriptome assemblies for each species often possess better assembly quality when compared to the combined assemblies by metrics such as N50 and mean contig length (Fig 2, Table 1). As a result of being made up of a subset of the total reads, the individual assemblies do not possess the breadth of the combined assemblies, with fewer contigs, especially at long contig length (e.g. greater than 1kb). As such, the staged transcriptomes were used solely for comparison of expression levels across time, while the combined assemblies were used for gene family analyses. We emphasise that no technical replicate was performed for these comparisons, and any conclusions drawn from them should hold this consideration in regard. With this limitation in mind, we carried out differential expression analyses and observed broad trends in expression, as can be seen in Fig 3.

**Figure 3:**
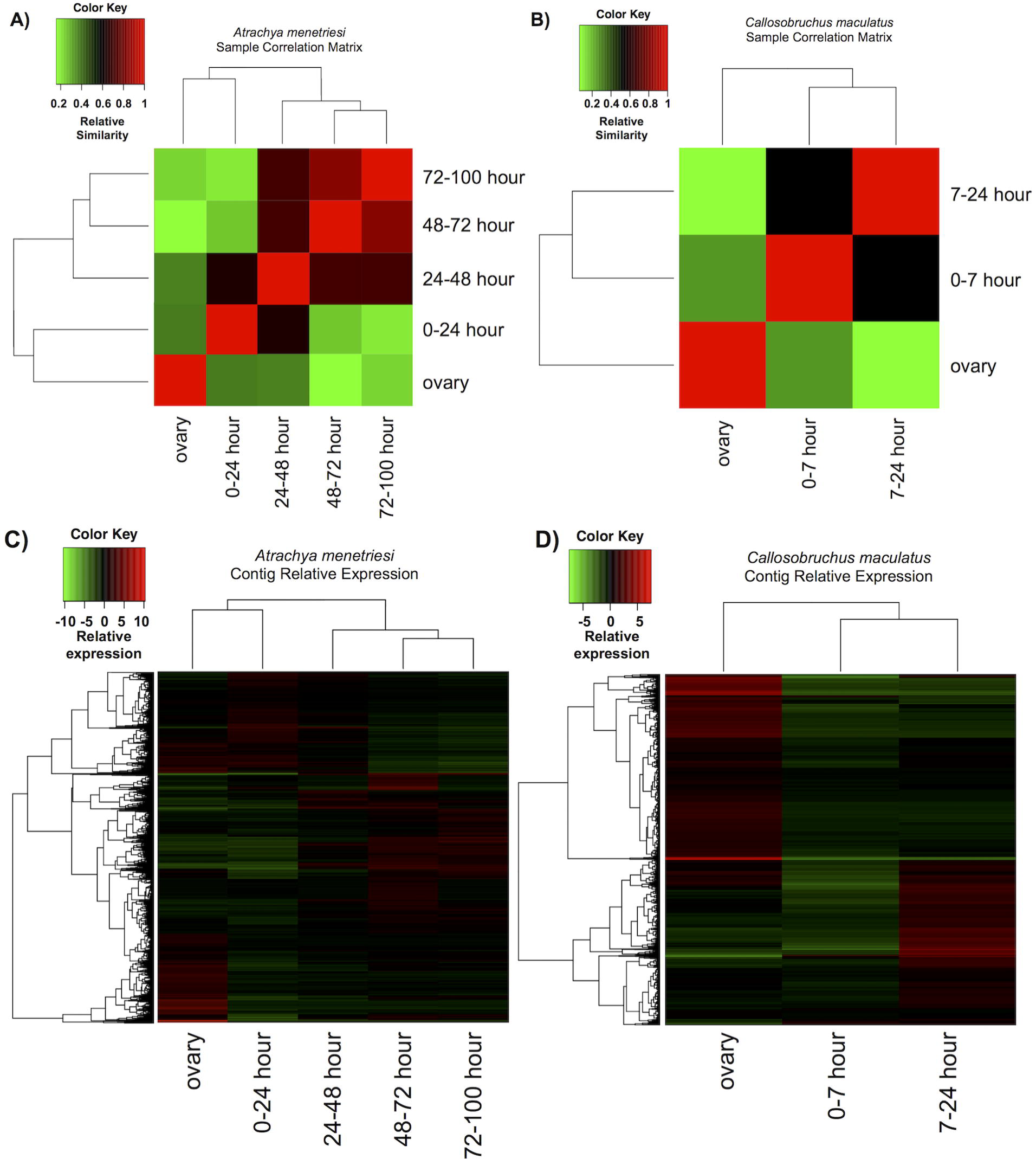
Overview of results of differential expression analysis performed by RSEM within the Trinity framework. A) and B) show the sample correlation matrix for *A. menetriesi* and *C. maculatus* respectively. C) and D) show relative expression of each differentially expressed contig, considered as isoforms, across time.

First, we generated matrices from general comparison between time points in order to find the most similar samples (Fig 3 A,B). Generally, these results are congruent with steady changes in gene expression across the course of development, with most time points being most similar to those immediately preceding and following them. However, a split can be seen in *A. menetriesi* (Fig 3 A,B) between the ovary and 0-24 hour samples and all others, with the three later samples resembling each other more closely than the 0-24 hour dataset resembles the 24-48 hour sample. This could be due to the maternal-zygotic transition, which will occur at some point in this time frame. This cannot be seen for *C. maculatus*, and could be due to more admixture of RNA within the last 7-24 hour sample. Focused analysis on when the maternal-zygotic transition occurs in each species is required to resolve this question.

Next, we clustered the results of our differential expression calculations (Fig 3 C,D). The results shown are those with RSEM considering each isoform separately (rather than taking into account clustering into genes performed by Trinity). Numerous up- and down-regulated contigs can be seen at each time point, with some time points more obviously possessing or lacking a subset of genes found in the combined transcriptomes.

While replicates have not been performed and we have not analysed up and down regulated transcripts in detail, these data are available to download from Additional Files S6 and S7 attached to this document online. These data will act as a good initial guide for those interested in tracking differential expression of specific genes across development and will be useful for hypothesis building.

### Basic Gene Annotation

To gain an understanding of the depth of coverage of our datasets we used the BUSCO library of well-annotated genes [48], which are known to be highly conserved in single copy across the Metazoa, as a basis for comparison with our combined, assembled transcriptomes. Of the BUSCO set of 843 metazoan orthologs, the *A. menetriesi* assembly possesses 801 (95%) complete (of which 225, 26%, appear to be duplicated), 16 fragmented (1.8%) and 26 missing genes (3.0%), for a total recovery of 97% of the BUSCO dataset. The *C. maculatus* assembly contains 815 (96%) complete (with 202, 23%, appear to be duplicated), 13 (1.5%) fragmented and 15 (1.7%) missing genes (98.3% recovery). This extremely high level of recovery gives us confidence that at least all housekeeping genes expected to be present in these species are found in our datasets, which strongly suggests that these transcriptomes contain the vast majority of the gene cassette of these species.

The number of potentially duplicated genes in our BUSCO analyses likely reflects the construction of our transcriptomes from mixed RNA samples, with the allelic variation that this implies. Discerning true duplicates from allelic and splice variant data is largely contingent on the availability of well-assembled genomes. However, the recovery of these putative duplicates in our assemblies underlines that our RNA sequencing and assembly was of good depth and quality (respectively). With this information in-hand, a range of investigations will be made possible, particularly into the regulation and expression of developmentally important genes.

A further understanding of the content of our assemblies was gained from Blast2GO analysis [49,50]. Genes were annotated using b2gpipe, on the basis of BLASTx (*E* value cutoff, 10^−3^) comparisons made against the nr database as downloaded on the 26 January 2015. Of 228,096 (*A. menetriesi)* and 128,837 (*C. maculatus)* contigs in each assembly, 78,879 (34.6%) and 41,744 (32.4%) possessed a hit in the nr databases above the threshold. After further annotation with ANNEX and Interproscan, a total of 36,315 (15.9%) and 13,096 (10.2%) contigs were assigned to one or more GO categories. These numbers, while only a fraction of the total number of contigs within our transcriptomes, more closely reflect the expected eukaryotic protein complement in number. Blast2GO annotations are given in Additional Files S2 and S3

Fig 4A shows the distribution of species best hit by BLASTx comparison of contigs from *A. mentriesi and C. maculatus* with the nr database. For both species, *T. castaneum* is the best represented species - a reflection of both the phylogenetic position of this species and its well annotated genome. The fact that contigs in our transcriptomes match *T. castaneum* and other coleopteran and insect species more closely than those of other species suggests that gene orthology will be easy to assign in many cases, and that an abnormally high rate of molecular evolution is not observed in our species.

**Figure 4:**
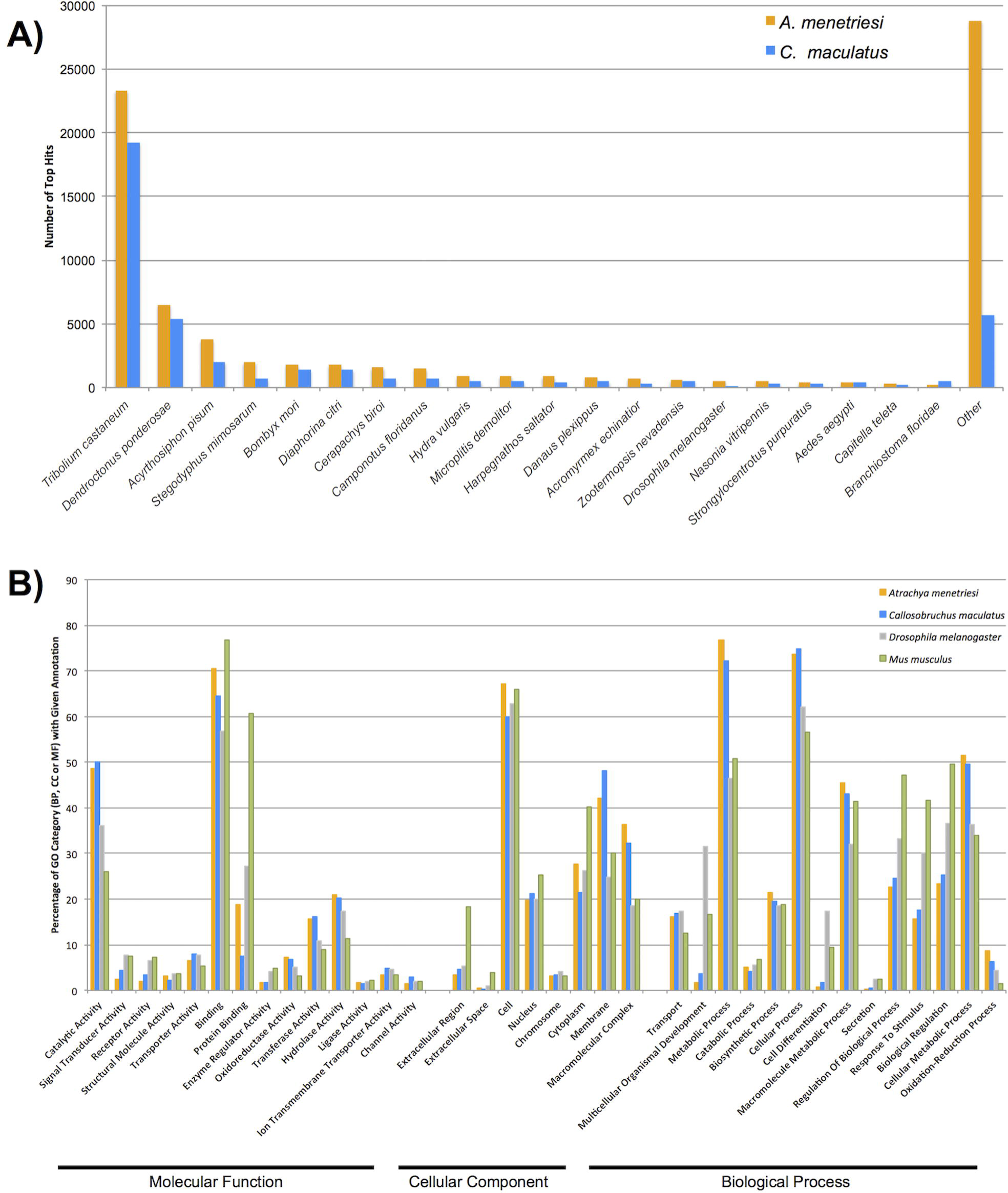
Blast2GO results A) Distribution of BLASTx best hits by species, showing metazoans only, for *A. menetriesi* in orange, *C. maculatus* in blue. B) Distribution of GO terms expressed as a percentage of annotated contigs which were assigned a term within each of the three (Molecular Function, Cellular Component, Biological Process) categories of GO ID.

The distribution of GO terms within our datasets are shown in Fig 4B, alongside those of the well-annotated *D. melanogaster* and *Mus musculus* proteomes. In general, our datasets resemble one another more than they mirror that of the two sequenced genomes noted. Our transcriptomic resources empirically seem under-represented relative to *D. melanogaster* and *M. musculus* in developmentally interesting categories such as ‘Protein Binding’ (Molecular Function), ‘Multicellular Organism Development’ and ‘Cell Differentiation’ (Biological Process). Given our results for more targeted investigations, found below, we feel this is likely a result of poor annotation of these by Blast2GO, rather than true absence from the transcriptome.

At a gross level, in the intracellular ‘Cellular Component’ GO categories, our data appears more similar to that seen in the two ‘model’ species than in Molecular Function or Biological Process categories, although this has not been tested statistically. This would suggest these well-conserved structural components were more readily assigned GO categories than the developmentally interesting categories listed above. While not all GO categories are as well-annotated by this process as may be desired, the broad classification of our data into a wide range of GO categories of all levels of GO distribution demonstrates that Blast2GO annotations of our data are a useful starting point for more focussed investigations and identification of specific genes and pathways of interest.

### Gene Family Recovery

To extend the semi-automated analyses presented above, we performed more targeted analysis of individual, developmentally important gene families. Both the Hox family of transcription factors and the TGF-β cassette were exceptionally well recovered in our dataset. The Hox genes, and in particular the ANTP HOXL class, which pattern the anterior-posterior body axis, are recovered almost in their entirety in both species when compared to well-catalogued databases (e.g. [51]). This can be seen in Fig 5, which shows the phylogenetic distribution of recovered ANTP HOXL class sequences from our transcriptomes alongside previously annotated members of this class.

**Figure 5:**
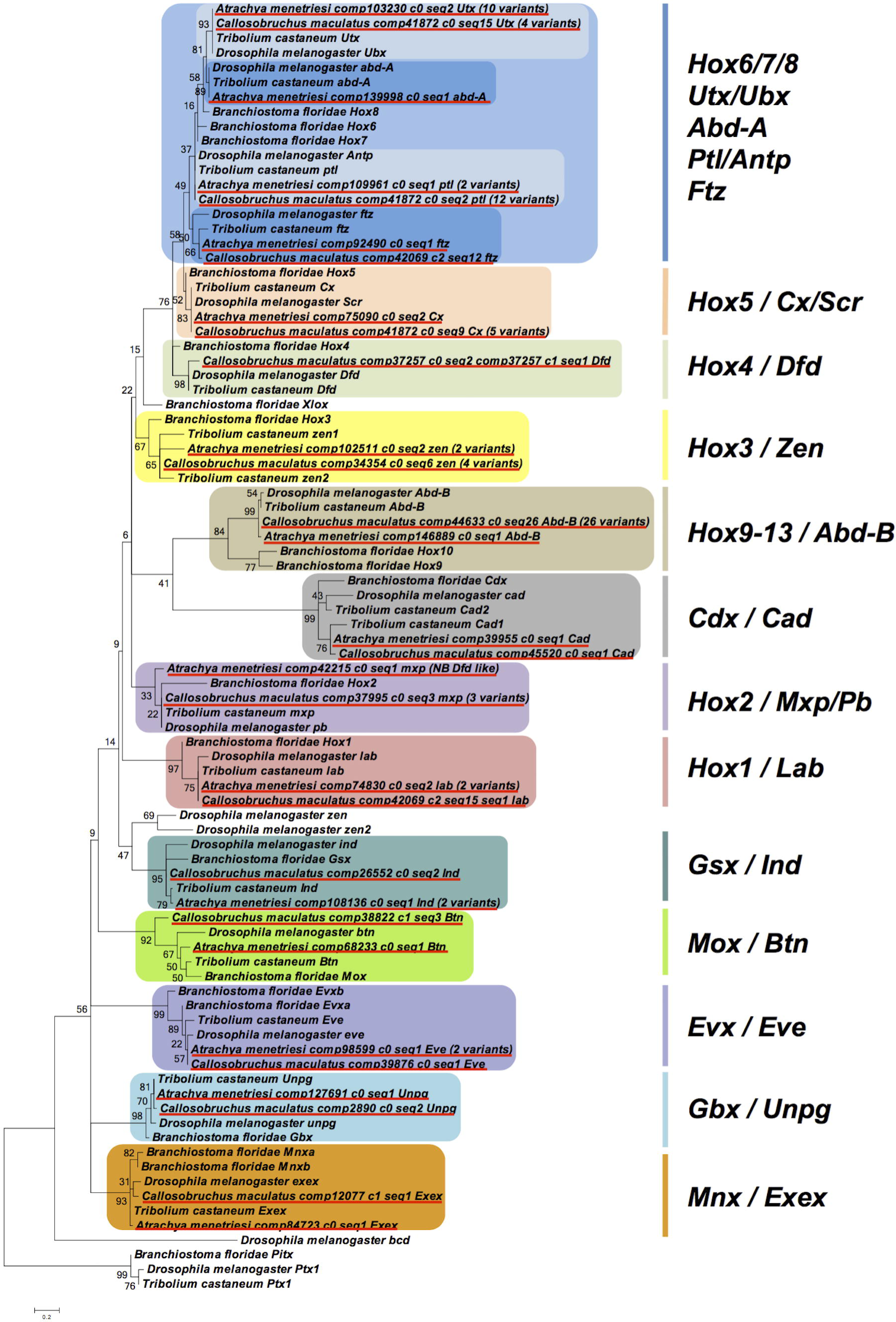
Phylogenetic inter-relationships of ANTP HOXL class genes, as reconstructed by MEGA 6 using the LG+Freqs model with 4 gamma categories and invariant sites, based on a 59 amino acid alignment spanning the homeodomain. Numbers at base of nodes represent bootstrap percentages of 1000 replicates. Scale bar at base of phylogeny gives substitutions per site at given unit distance. Red underline indicates *A. menetriesi* and *C. maculatus* sequences, coloured boxes are used to delineate known gene families (and in the case of Hox 6/7/8, a superfamily).

In *A. menetriesi,* sequences for all the ANTP HOXL class genes are recovered in our transcriptome, as can be seen in Fig 5, with sequences and alignments in Additional File S1. We note, however, that the *A. menetriesi Hox 2 / maxillopedia (mxp)/ proboscipedia (pb)* sequence bears some BLAST similarity with *Hox 4 / Dfd* sequences, while the putative *Hox 4 / Deformed (Dfd)* recovered does not include the homeodomain sequence - whether it has lost this crucial domain, or if this portion of the sequence is simply not recovered in our assembly is at present unknown. The putative *A. menetriesi Hox 4 / Dfd* sequence is given in Additional File S1, although it is not shown in the phylogeny in Fig 5 due to its truncated length.

While 2 (*A. menetriesi)* and 4 (*C. maculatus*) *Hox 3 / zerknüllt (zen)* variants are seen in our species, these are identical at the coding level, and therefore seem to be splice or allelic variants, rather than the two paralogous genes seen in *T. castaneum* [52]. Similarly, no evidence of the *caudal* (*cad*) duplication seen in *T. castaneum* [53] can be found in our transcriptomic assemblies, suggesting this is perhaps specific to the flour beetle lineage. Both *zen* and *caudal* are important embryonic patterning genes, and comparison of these genes in our two species and *Tribolium* would be an excellent situation in which to study how sub- and neo-functionalisation occurs. Likely allelic or splice variants are also observed for other HOXL genes in both *A. menetriesi* and *C. maculatus*. It should be noted that these could represent very recent duplications, or the effect of gene conversion, although this can only be tested fully with the advent of a complete genomic resource. No *Pdx/Xlox* gene was seen, adding further circumstantial evidence for the broad scale loss of this gene across the Insecta [54] and possibly the wider Arthropoda [55].

Our *C. maculatus* transcriptome also contains the full complement of ANTP HOXL genes, although, similar to the case of *Hox 4 / Dfd* in *A. menetriesi*, the 4 recovered *abdominal-A (abd-A)* homologs lack the whole homeodomain sequence, with several residues missing. These truncations excluded them from the phylogeny shown in Fig 5, but the sequences for these putative homologs are given in Additional File S1.

Even more so than in *A. menetriesi,* a remarkable diversity of potential splice/allelic variants are noted in *C. maculatus,* particularly for the *Hox 6/8* superfamily and *Hox 9-13* / *Abdominal-B* (*Abd-B*) gene family. Of the *Hox 6/8* gene superfamily, normally represented by four genes in *T. castaneum* (*prothoraxless* (*ptl)*, *fushi tarazu* (*ftz)*, *Ultrathorax* (*Utx)* and *abd-A*), 12 *ptl*, 1 *ftz*, 4 *Utx* and 4 *abd-A* representatives were found in our analysis. Furthermore, up to 26 different potential allelic or splice variants of abdB are recorded. As our transcriptome is made of mixed embryonic samples, it is perhaps not surprising that a diversity of putative splice/allelic variants are observed, but the excellent recovery of this data confirms the deep coverage provided by our sequencing and assembly given the coverage possessed by all isoforms.

The TGF-β cassettes of the insects have been very well described previously (e.g. [56,57]). Our datasets recover almost the full expected complement of the Coleoptera. A slight exception to this is *Activin (Act)*, a partial sequence of which is recovered for both species: a portion of the propeptide which does not span the mature signal peptide sequence. Whether this is a consequence of loss of the mature domain in these species or low levels of expression at the sampled timepoints remains to be established. A *BMPx* ortholog of clear homology to genes of that family can be found in *C. maculatus,* but has been excluded from the tree seen in Fig 6 as it is incomplete in length. Its sequence can be found in Additional File S1, and we have no doubt as to its identity due to high levels of sequence conservation between it and the *T. castaneum* and *A. menetriesi* orthologs of this gene.

**Figure 6:**
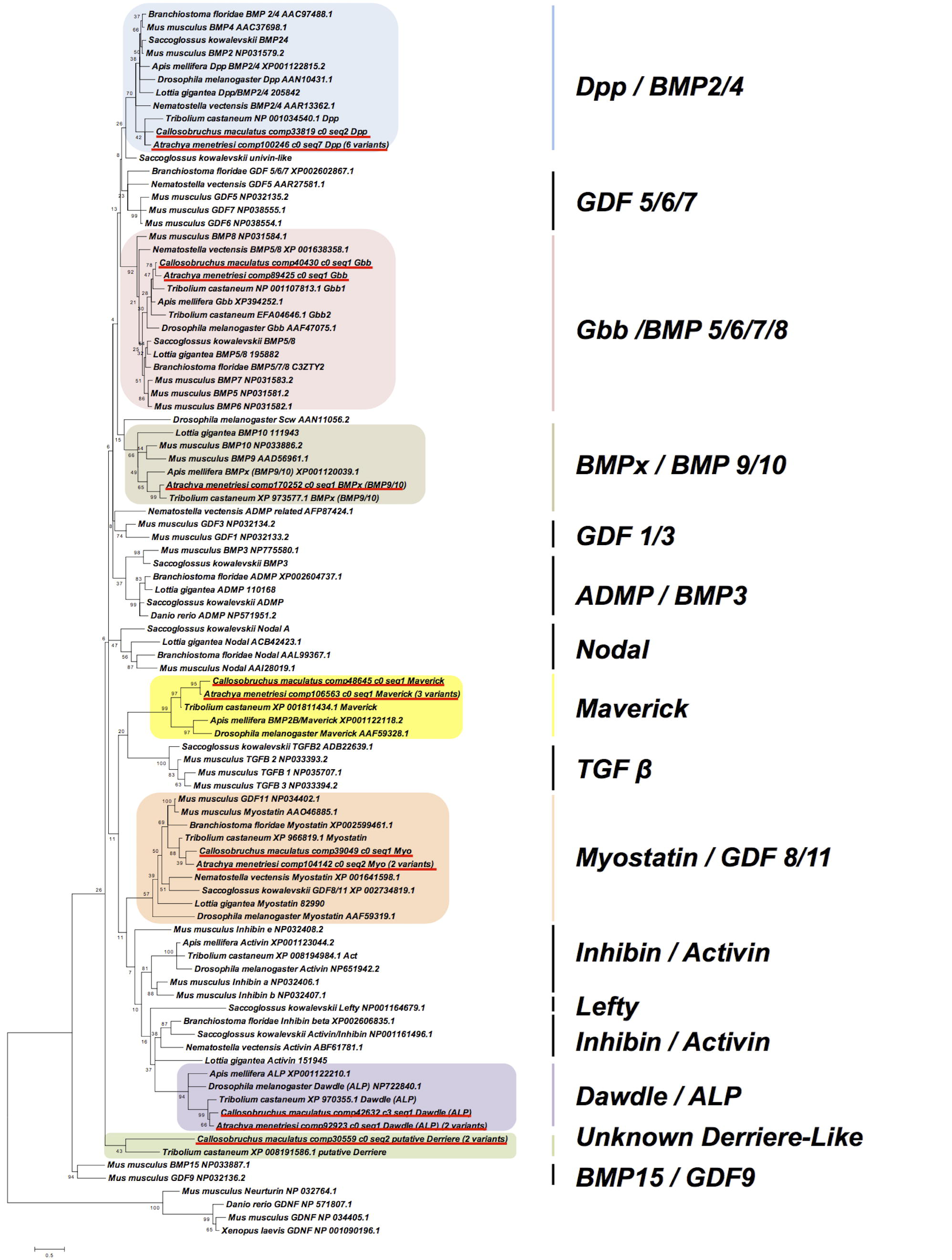
Maximum Likelihood Phylogeny of TGF-β ligands, as determined using MEGA under the LG+Freqs model with 4 gamma categories and invariant sites, on the basis of a 72 amino acid alignment of mature peptide sequences. The given scale depicts the number of substitutions per site per unit length. Bootstrap percentages (of 1000 replicates) are given at base of nodes. *A. menetriesi* and *C. maculatus* sequences are underlined in red. Coloured boxes represent known gene families with representatives in our transcriptomic resources, while all gene families, including those not found in our datasets, are indicated at right.

The *glass bottom boat (gbb)* duplication observed in *T. castaneum* cannot be found in our data, but we can recover a range of splice or allelic variants for other genes, especially in *A. menetriesi*. These do not differ in the protein coding regions, which leads us to suspect that these are not from gene duplications (unless the duplication(s) occurred very recently). The phylogeny shown in Fig 6 confirms the homology of all genes, and splice/allelic variant numbers observed are given there in brackets, with all sequences available in Additional File S1.

We also note the discovery of an additional putative TGF-β ligand in *C. maculatus*. This gene has been previously automatically annotated as *derriere* in *T. castaneum*, (XP_008191586.1) and if it is truly of this family, which is also known as *GDF1/3/Univin/Vg1*, this would be a surprise, as its presence outside the Deuterostomia is controversial [58]. If proof could be found for this being a *bona fide GDF1/3/Univin/Vg1*, the presence of this gene in more than one coleopteran could suggest that this might in fact be ancestrally present in all bilaterian species, but further investigation is warranted before strong conclusions can be drawn in this regard. We could gain no phylogenetic support for placing either of these beetle sequences in the GDF1/3 clade, and it may well be that these sequences instead represent a coleopteran novelty.

Our recovery of not only the full expected complements of these vital developmental genes, but also a remarkable diversity of alternative variants, demonstrates the depth of our assemblies as a resource, given the high coverage for each of these variants. Whether used as the basis for simple cloning or more sophisticated analysis of patterns of gene variation and diversification, these transcriptomes will be of wide utility to the field of coleopteran and insect developmental biology.

### Pathway Recovery

As well as examining specific gene families, we investigated a number of broader pathways commonly studied in insects [59]. This allows us to both note how well-recovered such pathways are in our species as a measure of transcriptome utility, as well as note interesting differences between these pathways in our species when compared to others. We did this using both automated (KEGG KAAS mapping) and manual (BLAST based) methods. Some representative results of KEGG KAAS mapping can be seen in Fig 7, and all KEGG annotations can be downloaded from Additional Files S4 and S5.

**Figure 7:**
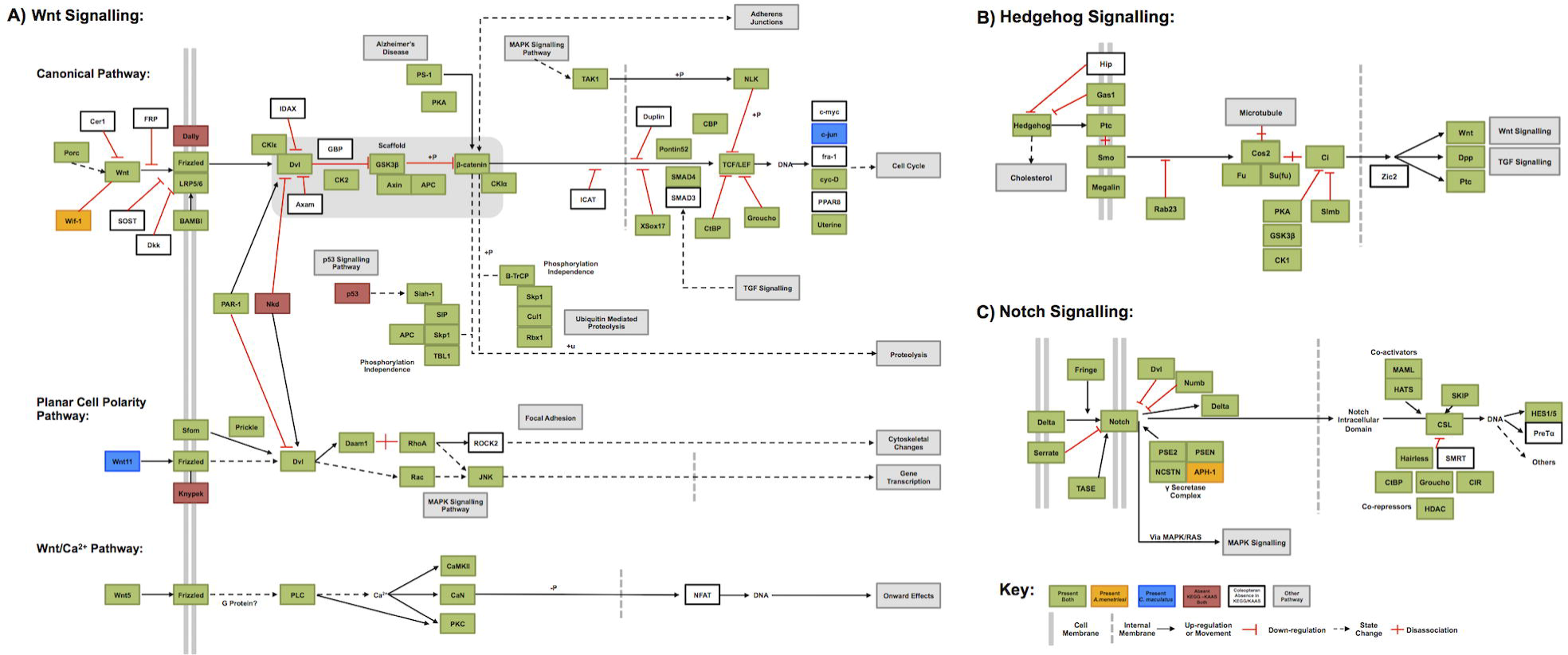
KEGG style pathway maps showing recovery in our transcriptome resources of A) the Wnt signalling pathway in canonical and non-canonical contexts, B) the Hedgehog signalling pathway and C) the Notch signalling pathway. Coloration of genes indicates presence, absence or ancestral absence from the Coleoptera as detailed in the key, which also gives other information as noted. All genes automatically annotated by KEGG KAAS server, with the exception of *PAR-1,* which was manually annotated.

KEGG KAAS mapping uses BLAST results to annotate known pathways, and gives a rapid overview of the recovery of these. Here we have shown the well-known Wnt, Notch and Hedgehog pathways to indicate the depth of our transcriptomes, and show how they may be useful for future research. However, these maps often use terminology based on vertebrate nomenclature, and contain genes known to be absent from particular clades. We have therefore indicated in Fig 7 (using unshaded boxes as shown in the Key) genes that may be absent ancestrally in the Coleoptera, based on their absence from the *T. castaneum* pathway. Of genes expected to be present in the Coleoptera we find almost total recovery in our transcriptomes. In the three pathways examined, only three genes noted to be present in *T. castaneum* were noted as absent from both of our transcriptome datasets, all in the Wnt cascades (Fig 7A). The expected Hedgehog cassette was recovered *in toto* (Fig 7B) and in the Notch signalling cascade (Fig 7C), only APH-1 was noted to be absent, and only from the *C. maculatus* transcriptome. We must note that these may not be true absences - KEGG KAAS mapping is based on automatic BLAST assignation, and if these sequences are divergent in our transcriptomes they may have been missed by this analysis.

We also examined pathways manually, using reciprocal BLAST hits and closer manual investigation to confirm the identities of individual genes, the results of which can be seen in Table 2. The anterior-posterior patterning genes *cad* (mentioned earlier) and *hunchback (hb)* are present in both species. Of the germline establishment and localization genes examined, *nanos* was surprisingly absent in both species, while *bruno (bru;* also known as *arrest*), *exuperantia (exu)*, *tudor* (*tud;* 2 copies in *A. menetriesi)*, *oskar (osk)*, *vasa (vas)* and *valois (vls)* were present. Interestingly, *pumilio (pum)* is present in *C. maculatus* in single copy (although it is divided across two contigs), while *A. menetriesi* possesses a total of seven copies. The different *A. menetriesi pum* copies varied both at the nucleotide level and in their amino acid sequences, strongly suggesting that they are in fact paralogs. In depth analysis of these genes is required to uncover why they have undergone several rounds of duplication. Orthologs of the *Drosophila* gene *swallow (swa)* could not be found in either of our transcriptome resources, nor is it present in several other insects (data not shown) and we suggest it may therefore be a schizophoran novelty.

**Table 2:**
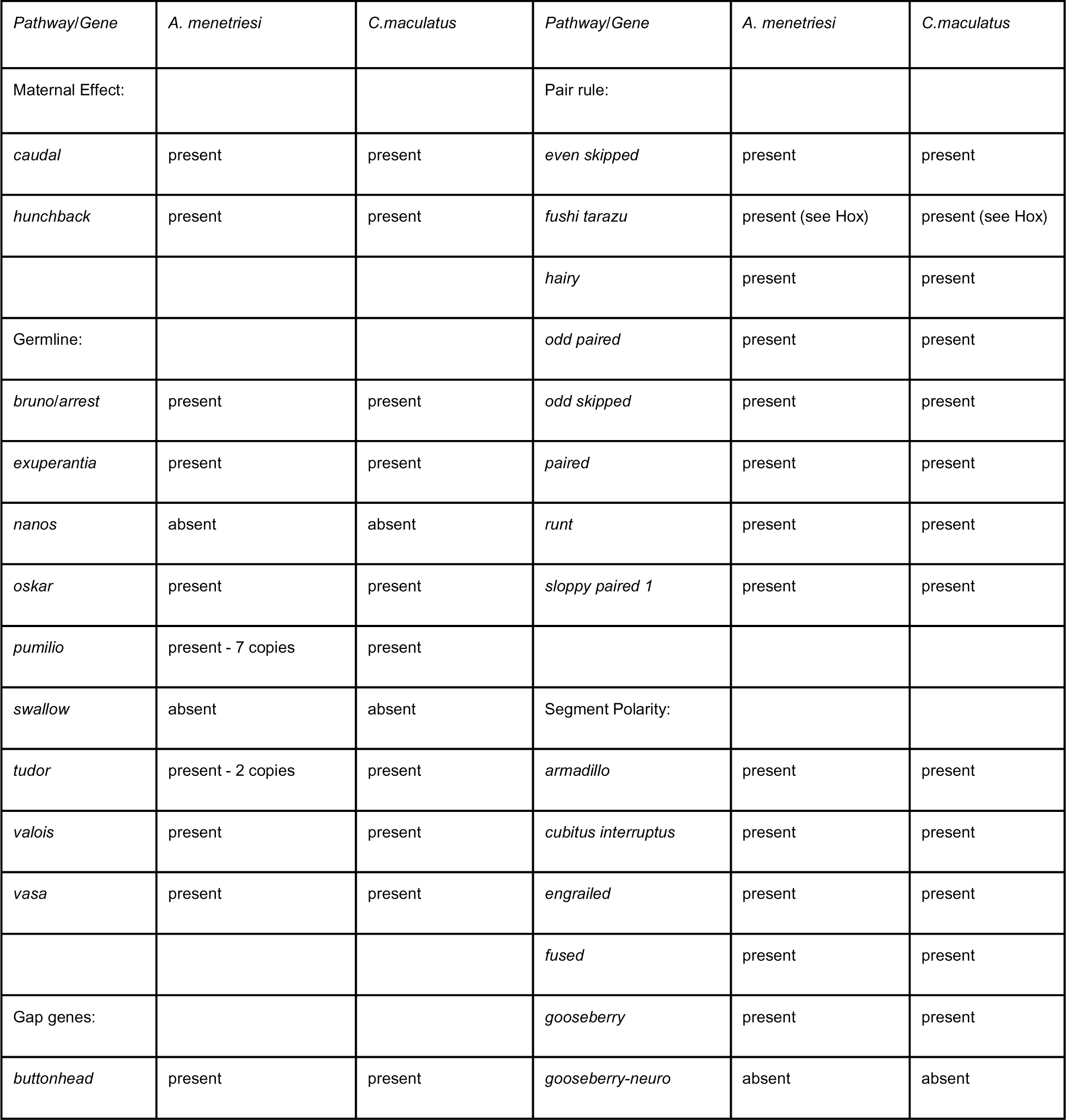
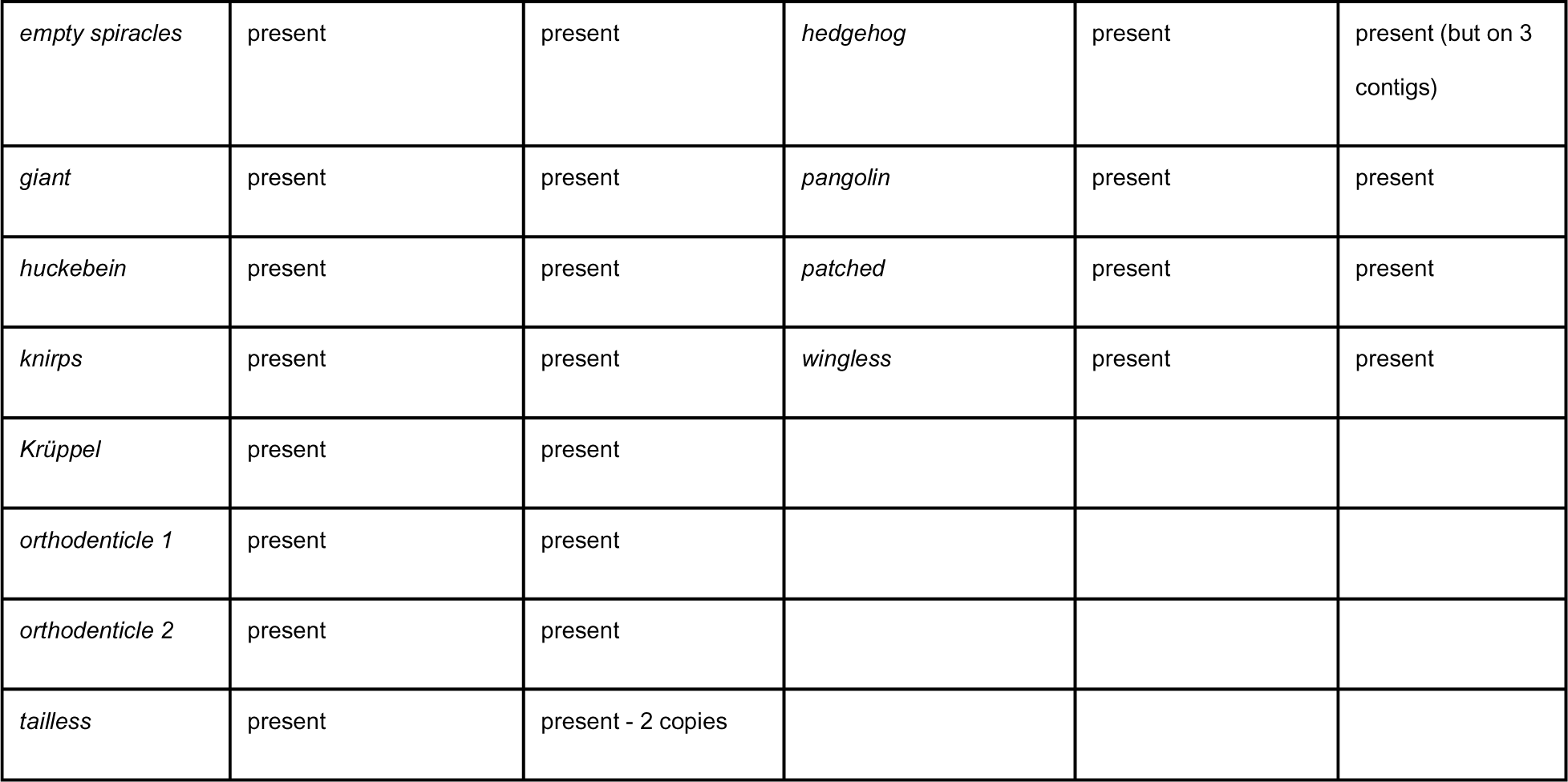
Genes identified by manual annotation

| *Pathway/Gene* | *A. menetriesi* | *C. maculatus* | *Pathway/Gene* | *A. menetriesi* | *C. maculatus* |
| --- | --- | --- | --- | --- | --- |
| Maternal Effect: |  |  | Pair rule: |  |  |
| *caudal* | present | present | *even skipped* | present | present |
| *hunchback* | present | present | *fushi tarazu* | present (see Hox) | present (see Hox) |
|  |  |  | *hairy* | present | present |
| Germline: |  |  | *odd paired* | present | present |
| *bruno/arrest* | present | present | *odd skipped* | present | present |
| *exuperantia* | present | present | *paired* | present | present |
| *nanos* | absent | absent | *runt* | present | present |
| *oskar* | present | present | *sloppy paired 1* | present | present |
| *pumilio* | present - 7 copies | present |  |  |  |
| *swallow* | absent | absent | Segment Polarity: |  |  |
| *tudor* | present - 2 copies | present | *armadillo* | present | present |
| *valois* | present | present | *cubitus interruptus* | present | present |
| *vasa* | present | present | *engrailed* | present | present |
|  |  |  | *fused* | present | present |
| Gap genes: |  |  | *gooseberry* | present | present |
| *buttonhead* | present | present | *gooseberry-neuro* | absent | absent |
| *empty spiracles* | present | present | *hedgehog* | present | present (but on 3 contigs) |
| *giant* | present | present | *pangolin* | present | present |
| *huckebein* | present | present | *patched* | present | present |
| *knirps* | present | present | *wingless* | present | present |
| *Krüppel* | present | present |  |  |  |
| *orthodenticle 1* | present | present |  |  |  |
| *orthodenticle 2* | present | present |  |  |  |
| *tailless* | present | present - 2 copies |  |  |  |

Canonical gap genes *Krüppel (Kr)*, *knirps (kni)*, *giant (gt)*, *huckebein (hkb)*, *tailless* (*tll;* 2 paralogs in *C. maculatus*), *buttonhead (btd)*, *empty spiracles (ems)* and both *orthodenticle* orthologs (*Otd* and *Otd2*) were recovered in both species examined here. The *C. maculatus* paralogs of *tll* exhibited differences at both the nucleotide and amino acid level along their entire lengths (data not shown) confirming that they are paralogs. Given the important embryonic role of *tailless* in other insects (for example [60]), this duplication would be excellent opportunity to study gene duplication and evolution. The pair rule genes e*ven skipped (eve), hairy (h), fushi tarazu (ftz), odd paired (opa), odd skipped (odd), paired (prd), runt (run)* and *sloppy paired 1 (slp1)* were present in single copy. The segment polarity genes were also present in both species, with the notable absence of *gooseberry-neuro (gsb-n)* from our datasets. The genes *armadillo (arm), cubitus interruptus (ci), engrailed (en), fused (fu), gooseberry (gsb), hedgehog (hh), pangolin (pan), patched (ptc)* and *wingless (wg)* were all present, and their sequences can be found in Additional File S1.

All of these pathways are commonly studied in insects, and the annotations provided here, along with preliminary timed expression data, will provide a basis for a wide range of targeted investigations into the embryonic development of these two species, and how these pathways have changed over the course of evolution. Furthermore, the excellent recovery of these pathways by both automated (KEGG-KAAS) and manual annotation gives us high confidence in the completeness of our transcriptomic resources. This confirms the results of our BUSCO analysis, and our datasets are therefore likely to contain the vast majority of transcribed genes in these two species, with only lowly expressed and temporally restricted genes absent from these transcriptome resources.

## Conclusions

Our production of deep transcriptomic sequence data for *A. menetriesi* and *C. maculatus* will assist in the inference of character gain and loss across the Coleoptera, aid in future phylogenetic efforts, and allow a range of investigations into the embryonic development of these species at the molecular level. The status of these organisms as common agricultural pests also suggests that such resources may allow targeted control mechanisms to be developed for these species. This data will be another key building block in our understanding of the transcriptomic basis to embryological development, and provide a window into the basic biology of the most successful clade of animals.

## Acknowledgements

The authors would like to thank the members of their laboratories for their support and discussions. In addition, we thank Dr Y Ando for providing *A. menetriesi* eggs and advice on establishing cultures, Dr J Savard for providing *C. maculatus*, and Dr F Marletaz for help in running BUSCO analyses. The efforts of editors and reviewers in considering this manuscript are gratefully acknowledged.

## Availability of data and materials

The datasets supporting the conclusions of this article are available in the NCBI SRA repository [Bioproject Accession numbers: PRJNA293391 and PRJNA293393] and in the Figshare repository [DOIs: 10.6084/m9.figshare.2056464.v2, 10.6084/m9.figshare.2056467.v2].

## Supporting Information Captions

Additional File S1: Sequences of all genes referred to in text, and alignments used in phylogenetic analyses (.xls)

Additional File S2: Annotations of transcriptome (Blast2GO.annot file) *A. menetriesi*

Additional File S3: Annotations of transcriptome (Blast2GO.annot file) *C. maculatus*

Additional File S4: KEGG data, *A. menetriesi*

Additional File S5: KEGG data, *C. maculatus*

Additional File S6: Comparative expression data, *A. menetriesi*

Additional File S7: Comparative expression data, *C. maculatus*

